# The STING pathway does not contribute to *Pink1/parkin* phenotypes in *Drosophila*

**DOI:** 10.1101/806265

**Authors:** Juliette J. Lee, Simonetta Andreazza, Alexander J. Whitworth

## Abstract

Mutations in *PINK1* and *Parkin/PRKN* cause the degeneration of dopaminergic neurons in familial forms of Parkinson’s disease but the precise pathogenic mechanisms are unknown. The PINK1/Parkin pathway has been described to play a central role in mitochondrial homeostasis by signaling the targeted destruction of damaged mitochondria, however, how disrupting this process leads to neuronal death until recently was unclear. An elegant study in mice revealed that the loss of *Pink1* or *Prkn* coupled with an additional mitochondrial stress resulted in the aberrant activation of the innate immune signaling, mediated via the cGAS/STING pathway, causing degeneration of dopaminergic neurons and motor impairment. Genetic knockout of *Sting* was sufficient to completely prevent neurodegeneration and accompanying motor deficits. To determine whether Sting plays a conserved role in *Pink1/parkin* related pathology, we tested for genetic interactions between *Sting* and *Pink1/parkin* in *Drosophila*. Surprisingly, we found that loss of *Sting*, or its downstream effector *Relish*, was insufficient to suppress the behavioral deficits or mitochondria disruption in the *Pink1/parkin* mutants. Thus, we conclude that phenotypes associated with loss of *Pink1/parkin* are not universally due to aberrant activation of the STING pathway.

## Introduction

Loss of function mutations in *PINK1* and *PRKN* cause familial parkinsonism, an incurable neurodegenerative disorder predominantly associated with the progressive loss of dopaminergic neurons in substantia nigra leading to loss of motor control. *PRKN* encodes a cytosolic ubiquitin E3 ligase, Parkin, and *PINK1* encodes a mitochondrially targeted kinase. Extensive evidence shows that they cooperate in signaling the targeted autophagic destruction of damaged mitochondria (mitophagy) as part of a homeostatic mitochondrial quality control process^1,2^.

Mitochondria are essential organelles that perform many critical metabolic functions but are also a major source of damaging reactive oxygen species (ROS) and harbor pro-apoptotic factors. Hence, multiple homeostatic processes, such as mitophagy, operate to maintain mitochondrial integrity and prevent potentially catastrophic consequences. Such homeostatic mechanisms are particularly important for post-mitotic, energetically demanding tissues such as nerves and muscles.

The molecular details of PINK1/Parkin-induced mitophagy are well characterized in cultured cells, however, relatively little is known about mitophagy under physiological conditions *in vivo*^3–5^. Nevertheless, several studies provide evidence consistent with PINK1 and Parkin acting to remove mitochondrial damage *in vivo*. One study used a mass spectrometry-based analysis of mitochondrial protein turnover in *Drosophila*^6^, which revealed that fly PINK1 and Parkin selectively affect the degradation of certain mitochondrial proteins under physiological conditions. Another found that loss of *Prkn* in mice, which alone has very little phenotype^7,8^, exacerbated the phenotypic effects of a mitochondrial DNA mutator strain, provoking loss of dopaminergic neurons and motor deficits^9^.

Importantly, a subsequent study shed light on the mechanism by which loss of *Pink1/Prkn* leads to neurodegeneration in the presence of mtDNA mutations, or upon exposure to exhaustive exercise, as chronic or acute mitochondrial stresses, respectively^10^. This demonstrated that in the absence of *Pink1/Prkn* these mitochondrial stresses cause an aberrant inflammatory response mediated by the STING pathway, likely via the release of mtDNA into the cytosol. Consequently, loss of *STING* completely prevented the inflammatory response and resulting neurodegeneration and locomotor phenotypes^10^. These results strongly implicate the induction of STING-mediated inflammation in the pathogenic cause of Parkinson’s disease.

The recently identified *Drosophila Sting* ortholog has been shown to bind to cyclic-dinucleotides and trigger an immune response to bacterial and viral infection^11–13^, mediated by the IMD pathway and the transcription factor Relish (homologous to NF-κB). Consequently, *Drosophila* mutant for *Sting* showed a reduced survival upon infection. Interestingly, while aberrant activation of the IMD-Relish pathway has been shown to cause neurodegeneration and shortened lifespan in *Drosophila*^14^, transcriptional profiling has shown that innate immune signaling pathways are ectopically active in *Drosophila parkin* mutants^15^.

The *Drosophila* models have been highly informative for interrogating the physiological role of PINK1/Parkin, primarily because of the robust neuromuscular phenotypes associated with loss of the *Pink1/parkin* orthologs^16–19^. Therefore, we sought to determine whether aberrant activation of the Sting-Relish immune signaling cascade may be responsible for the neuromuscular degeneration phenotypes observed in *Drosophila Pink1/parkin* mutants. Surprisingly, we found that loss of *Sting* or *Relish* had no suppressing effect on the locomotor deficits or mitochondrial disruption in *Pink1* or *parkin* mutants. Moreover, *Sting* knockout did not affect the behavioral phenotypes associated with a fly mtDNA mutator model, nor the combined effect of mtDNA mutations in a *parkin* background. Hence, the central role of Sting in the induction of *Pink1/parkin* mutant phenotypes is not conserved in *Drosophila*.

## Results

*Drosophila Sting* mutants have recently been generated and, consistent with Sting’s role in triggering an innate immune response, shown to be more susceptible to infection. As other organismal phenotypes were not reported^11–13^ we first assessed whether loss of *Sting* may induce additional phenotypes associated with the neuromuscular system that might confound further genetic interaction analysis. To this end, we examined the motor behavior and muscle integrity in *Sting* loss of function conditions. First, we assessed the impact of RNAi-induced loss of function using the ubiquitous driver *da-GAL4.* A small impact on climbing ability in young flies was observed with one RNAi transgene, which was also seen in homozygous *Sting* null (*Sting*^ΔRG5^) mutants (Fig. 1A). Aged *Sting*-RNAi flies showed a consistent, modest impact on climbing ability, but this was not evident in *Sting* mutants (Fig. 1B). Microscopy analysis of muscle and mitochondrial integrity did not reveal any obvious disruption in *Sting* mutants (Fig.1C). Since loss of Sting did not appear to grossly affect neuromuscular integrity, we next assessed whether the activity of Sting contributed to the neuromuscular phenotypes in *Pink1/parkin* mutants.

**Figure 1.**
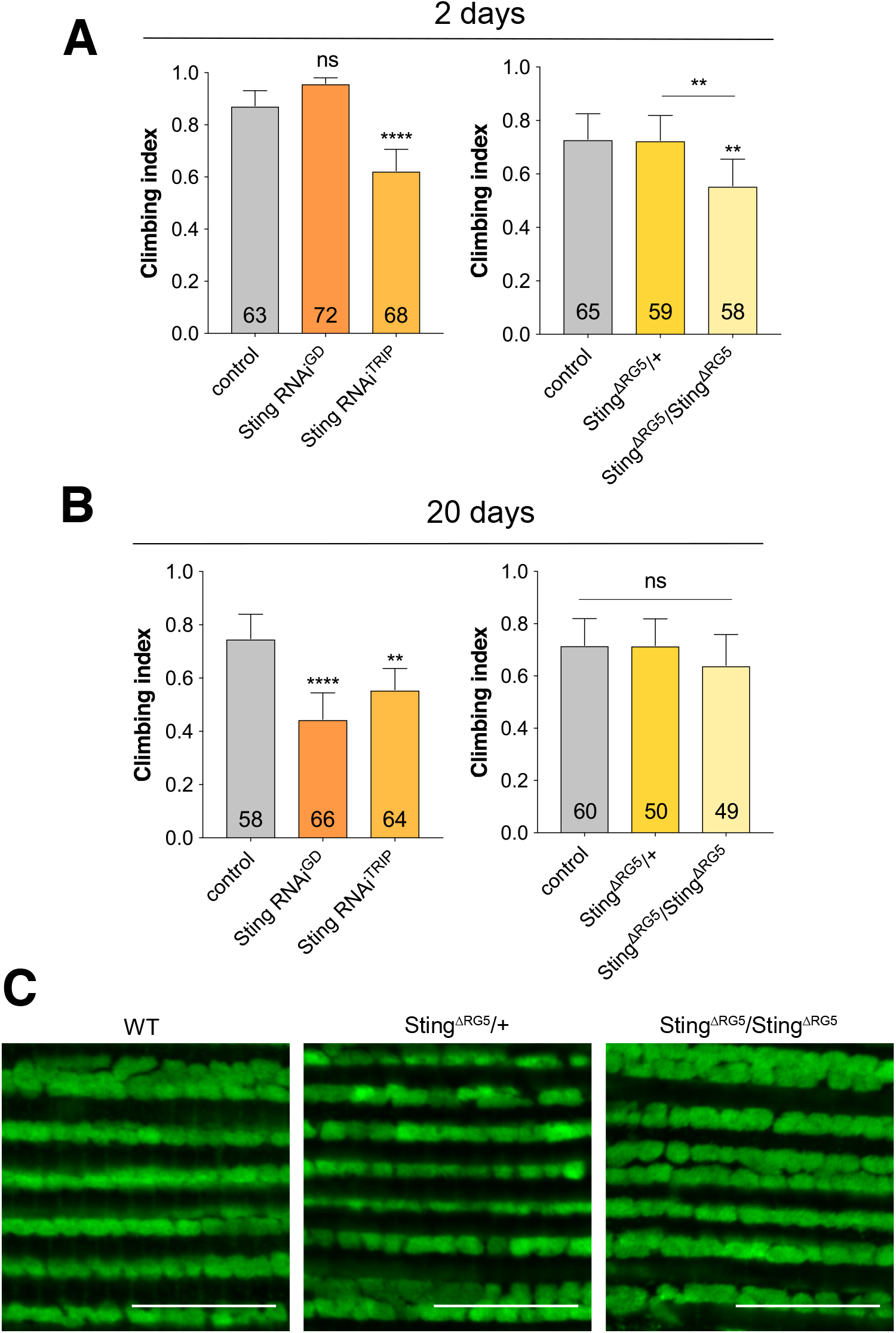
Loss of *Sting* has limited impact on neuromuscular phenotypes. Locomotor assays analyzing climbing ability (negative geotaxis) in (A) young and (B) older adult flies of control and *Sting* knockdown (RNAi) or null (*Sting*^ΔRG5^) mutants. Charts show mean ± 95% confidence interval (CI); number of animals analyzed is shown in each bar. Significance was measured by Kruskal-Wallis test with Dunn’s post hoc correction for multiple comparisons; ** p < 0.01, **** p <0.0001; ns, non-significant. Control genotype is *da-GAL4*/+. (C) Representative confocal microscopy analysis of mitochondria in flight muscles, immunostained with anti-ATP5A, in wild type (*w*^1118^) and *Sting* heterozygous and homozygous mutants. Scale bar = 10 µm.

Combining all the manipulations of *Sting* (two RNAi transgenes, heterozygous and homozygous null mutations) with *parkin* null mutants (*park*^25^), we did not observe any modification (suppression or enhancement) of the *parkin* mutants climbing defect (Fig. 2A). Similarly, the thoracic indentations typically observed in *park*^25^ flies due to the degeneration of the underlying musculature, was still present in the absence of *Sting* (Fig. 2B). Consistent with this, we did not observe any improvement of the tissue or mitochondrial integrity in the flight muscles of *parkin* mutants by removal of *Sting* (Fig. 2C).

**Figure 2.**
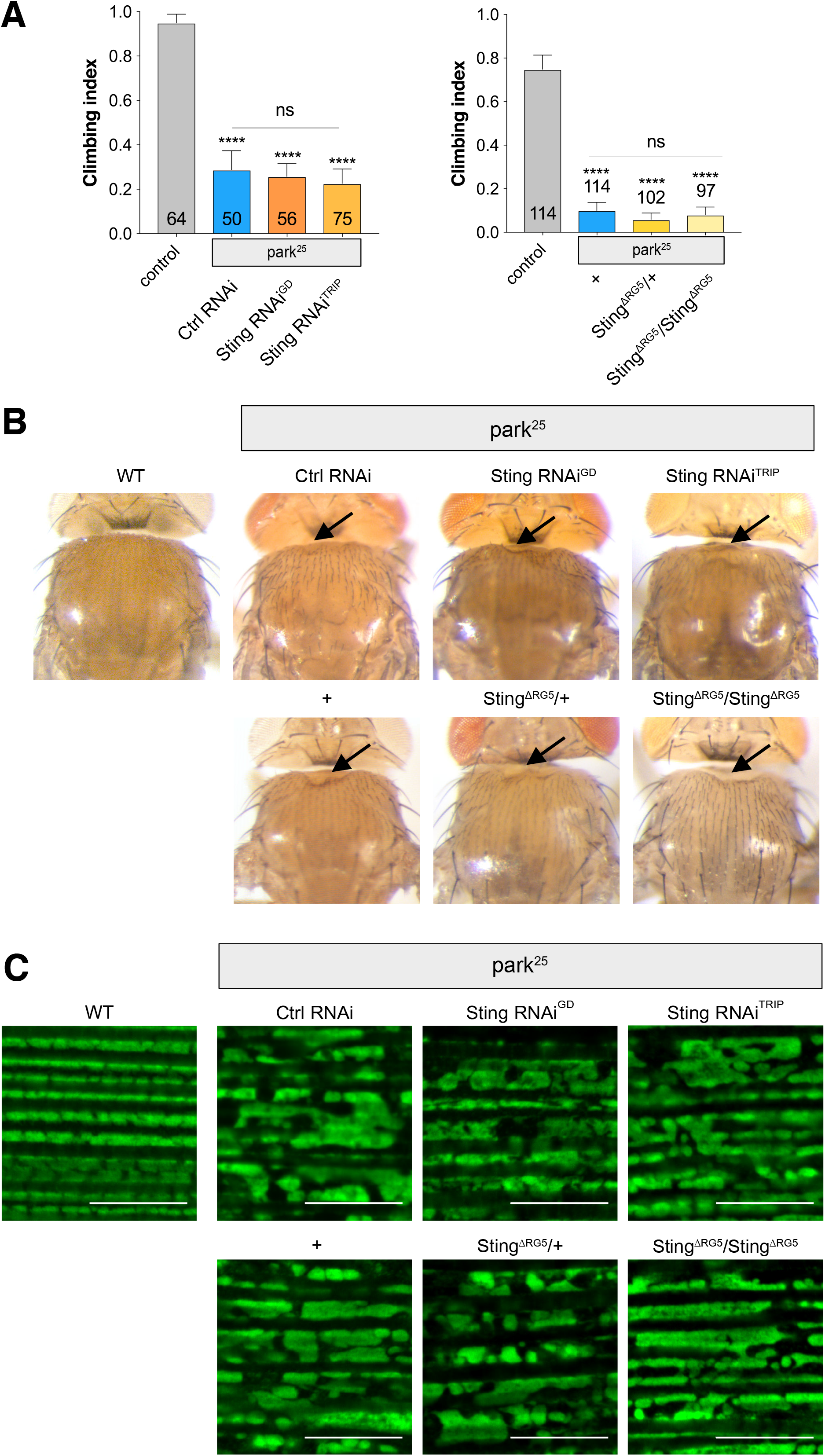
Loss of *Sting* does not modify *parkin* mutant phenotypes. (A) Analysis of locomotor (climbing) ability, (B) thoracic indentations, and (C) mitochondrial morphology in *park*^25^ mutants combined with *Sting* knockdown or null mutations. Charts show mean ± 95% confidence interval (CI); number of animals analyzed is shown in each bar. Statistical significance was measured by Kruskal-Wallis test with Dunn’s post hoc correction for multiple comparisons; **** p <0.0001; ns, non-significant. Confocal microscopy images show flight muscle mitochondria immunostained with anti-ATP5A. Scale bar = 10 µm. Control/WT genotype is *da-GAL4*/+ for climbing and *w*^1118^ for thoracic indentation and microscopy.

We next assessed the contribution of Sting function towards *Pink1* mutant (*Pink1*^B9^) phenotypes. Similar to *parkin* mutants, loss of *Sting* failed to modify the climbing defect (Fig. 3A), thoracic indentations (Fig. 3B) or disruption of flight muscle and mitochondrial integrity (Fig. 3C) observed in *Pink1*^B9^ flies. Taken together, these results indicate that Sting does not contribute to the neuromuscular phenotypes observed in *Pink1/parkin* mutants.

**Figure 3.**
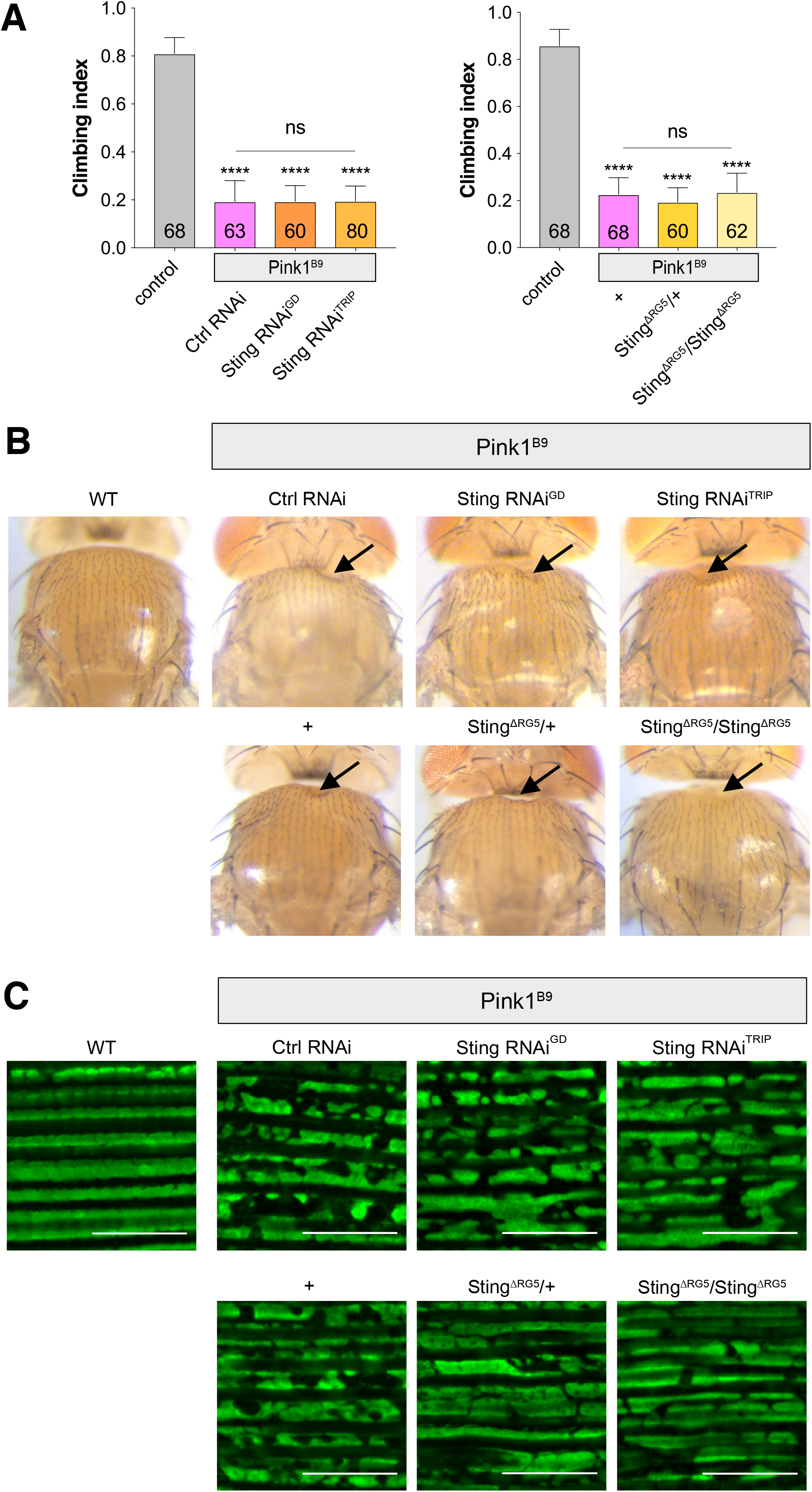
Loss of *Sting* does not modify *Pink1* mutant phenotypes. (A) Analysis of locomotor (climbing) ability, (B) thoracic indentations, and (C) mitochondrial morphology in *Pink1*^B9^ mutants combined with *Sting* knockdown or null mutations. Charts show mean ± 95% confidence interval (CI); number of animals analyzed is shown in each bar. Statistical significance was measured by Kruskal-Wallis test with Dunn’s post hoc correction for multiple comparisons; **** p <0.0001; ns, non-significant. Confocal microscopy images show flight muscle mitochondria immunostained with anti-ATP5A. Scale bar = 10 µm. Control/WT genotype is *da-GAL4*/+ for climbing and *w*^1118^ for thoracic indentation and microscopy.

Considering that loss of *STING* in mouse completely abrogated the *Pink1/Prkn*-associated neurodegeneration and motor phenotypes provoked by additional mitochondrial stresses, we were surprised by the lack of suppression of *Pink1/parkin* phenotypes in flies. Therefore, to further interrogate the potential contribution of this pathway to *Pink1/parkin* pathology, we also analyzed a downstream effector of the Sting-IMD pathway, the transcription factor Relish (Rel). While RNAi knockdown using two previously characterized transgenes^11,13^ elicited modest effect on climbing, *Rel* mutants (*Rel*^E20^) displayed a strong locomotor defect (Fig. 4A). However, analysis of flight muscles in these mutants did not reveal any major disruption of mitochondrial integrity (Fig. 4B).

**Figure 4.**
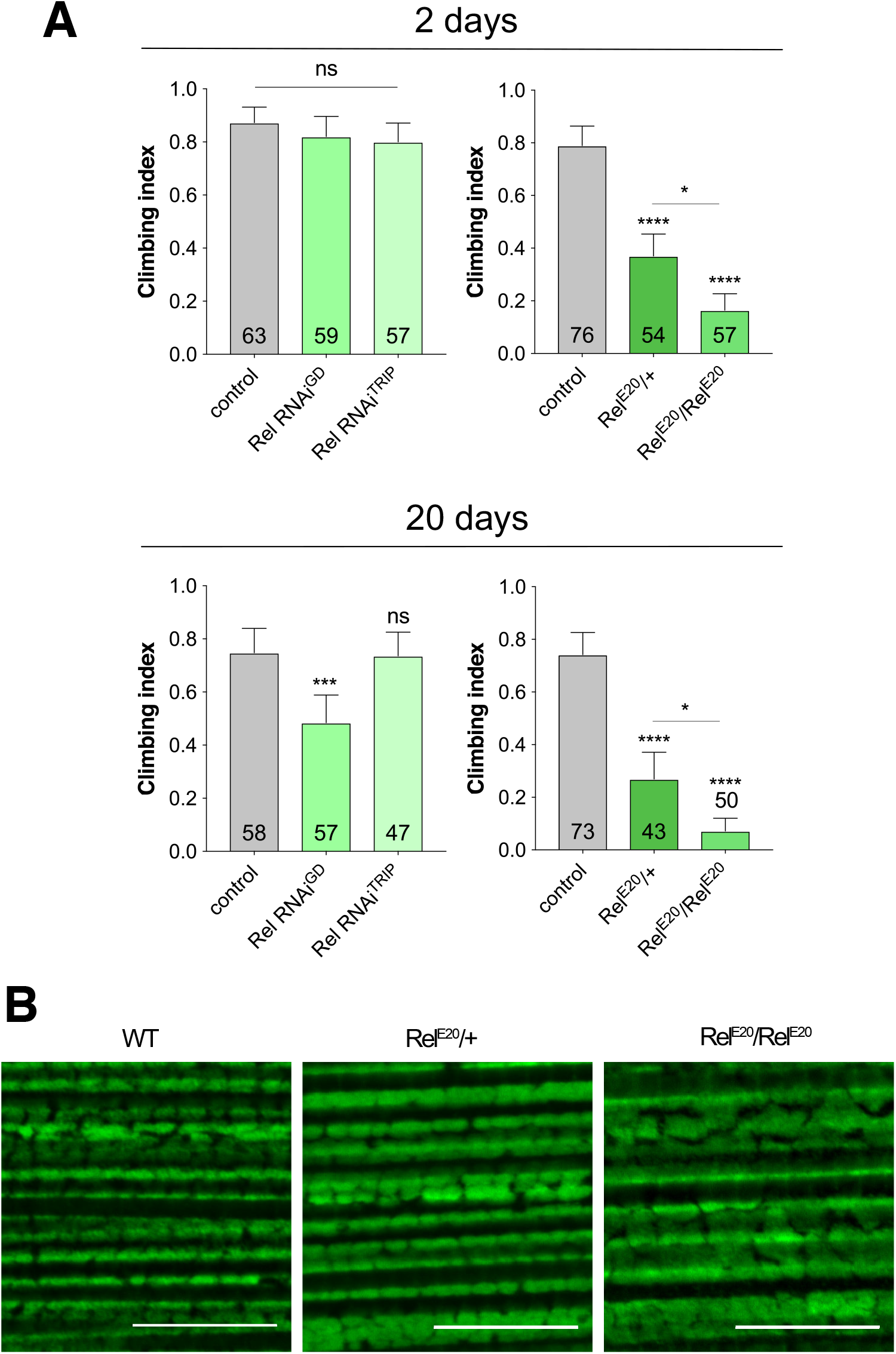
Loss of *Relish* causes mild locomotor deficits. Locomotor assays analyzing climbing ability in (A) young and (B) older adult flies of control and RNAi knockdown or *Relish* mutant (*Rel*^e20^). Charts show mean ± 95% confidence interval (CI); number of animals analyzed is shown in each bar. Statistical significance was measured by Kruskal-Wallis test with Dunn’s post hoc correction for multiple comparisons; * p < 0.05, *** p < 0.001, **** p <0.0001; ns, non-significant. Control genotype is *da-GAL4*/+. (C) Representative confocal microscopy analysis of mitochondria in flight muscles, immunostained with anti-ATP5A, in wild type (*w*^1118^) and *Relish* heterozygous and homozygous mutants. Scale bar = 10 µm.

Similar to the *Sting* manipulations, RNAi knockdown of *Rel* did not modify the climbing deficit of *parkin* or *Pink1* mutants (Fig. 5A), nor did it noticeably affect the mitochondrial integrity in flight muscles (Fig. 5B). Indeed, in contrast to expectation, genetic loss of *Rel* enhanced the *Pink1* locomotor defect (Fig. 5A), although the mitochondrial integrity was not noticeably worsened in *Rel*^E20^;*Pink1*^B9^ flies (Fig. 5B).

**Figure 5.**
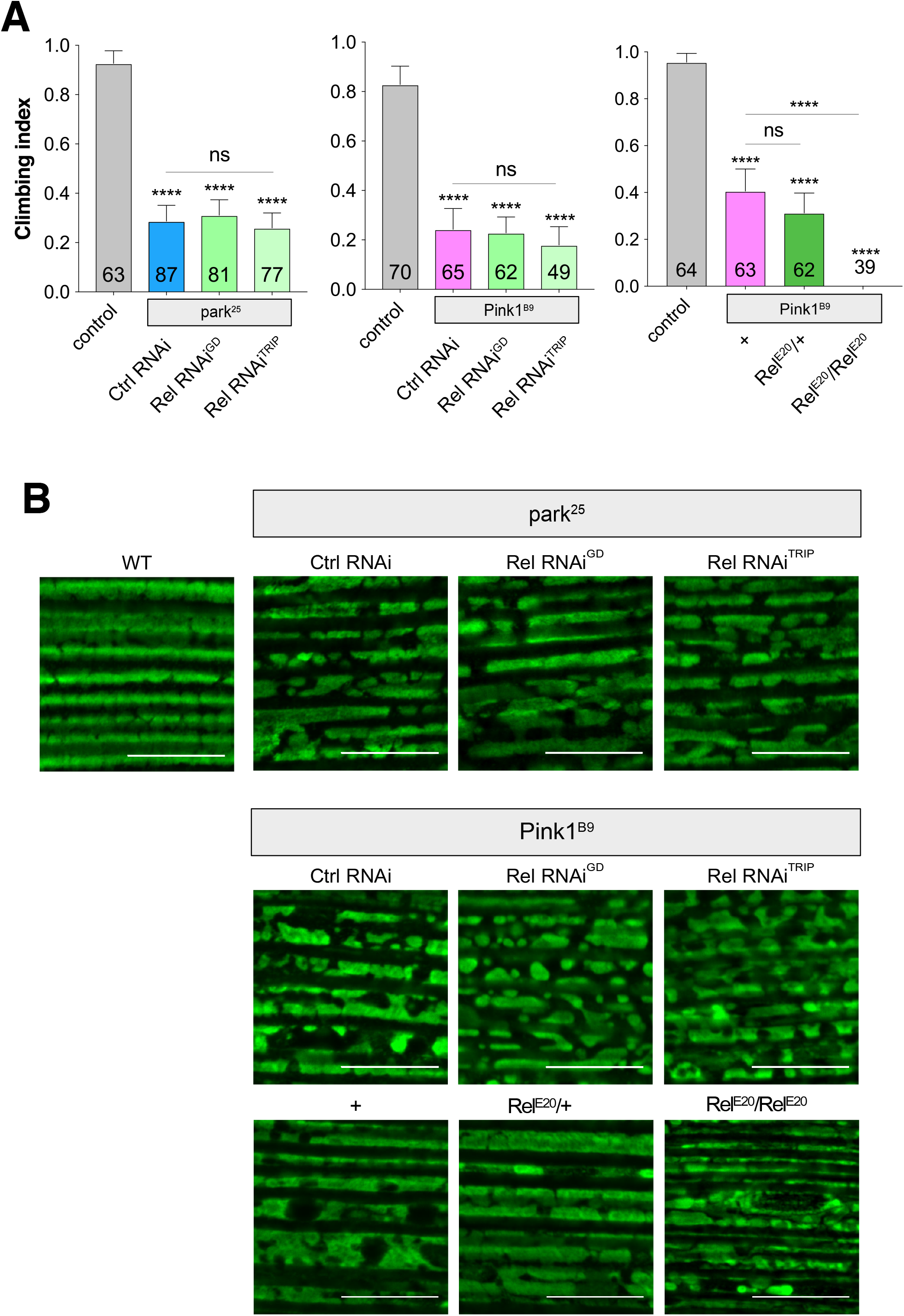
Loss of *Relish* does not rescue *Pink1* or *parkin* mutant phenotypes. (A) Analysis of locomotor (climbing) ability and (B) mitochondrial morphology in *park*^25^ or *Pink1*^B9^ mutants combined with *Relish* knockdown or null mutations. Charts show mean ± 95% confidence interval (CI); number of animals analyzed is shown in each bar. Statistical significance was measured by Kruskal-Wallis test with Dunn’s post hoc correction for multiple comparisons; **** p <0.0001; ns, non-significant. Confocal microscopy images show flight muscle mitochondria immunostained with anti-ATP5A. Scale bar = 10 µm. Control/WT genotype is *da-GAL4*/+ for climbing and *w*^1118^ for microscopy.

In a final effort to assess whether the *Drosophila* Pink1/parkin-Sting axis acts in an analogous fashion to mice, we sought to recapitulate the conditions assessed by Sliter et al.^10^ and test the role of Sting when an additional mitochondrial stress is combined with *parkin* loss-of-function. To do this, we used our previously established mtDNA mutator model (mito-APOBEC1), which generates high levels of deleterious mtDNA mutations in somatic tissues, disrupting mitochondrial function and causing motor defects and shortened lifespan^20^. Notably, the loss of *parkin* or *Sting* did not exacerbate the impact of mito-APOBEC1 alone on locomotor function (Fig. 6). Furthermore, the combination of the mtDNA mutator in a *parkin*;*Sting* double mutant background, in stark contrast to the results in mice^10^, enhanced the climbing deficit (Fig. 6).

**Figure 6.**
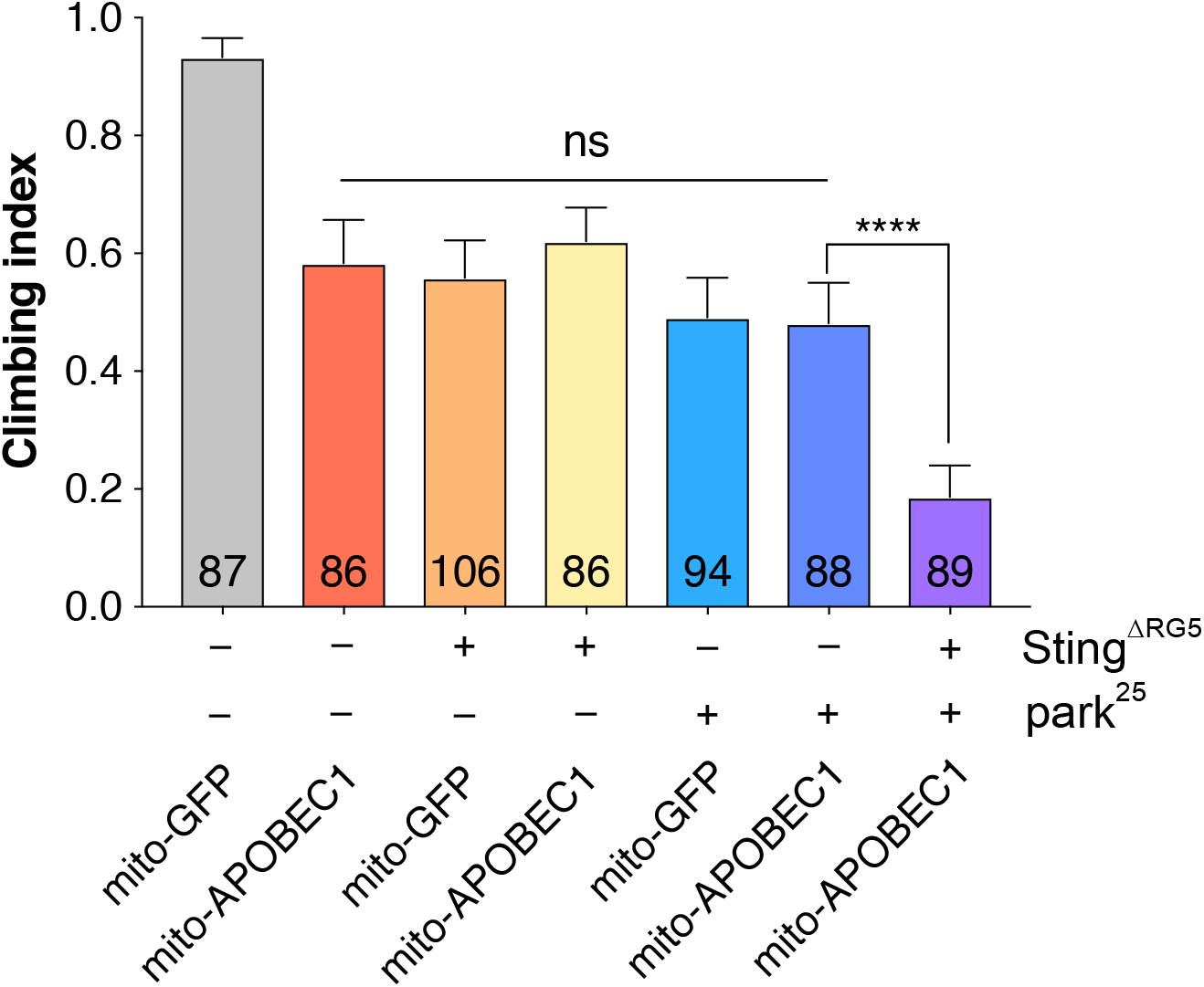
Loss of *Sting* does not ameliorate mtDNA *mutator*;*parkin* mutant combinations. Analysis of locomotor (climbing) ability in flies combining *mito-APOBEC1* mtDNA mutator expression with or without *parkin* and/or *Sting*. Transgene expression was driven by *da-GAL4*. *Mito-GFP* expression was used as a control. Charts show mean ± 95% confidence interval (CI); number of animals analyzed is shown in each bar. Statistical significance was measured by Kruskal-Wallis test with Dunn’s post hoc correction for multiple comparisons; **** p <0.0001; ns, non-significant.

Thus, together the above data suggest that the Sting pathway, although proposed to be mediating motor and neurodegenerative defects in *Prkn*^−/−^ mice, do not similarly contribute to the neuromuscular defects observed in *Pink1/parkin* mutant flies.

## Discussion

Understanding the pathogenic mechanisms by which loss of function mutations in PINK1 and Parkin lead to neurodegeneration in Parkinson’s disease is central to defining better disease-modifying therapies. While tremendous advances have been made in uncovering the molecular mechanisms of PINK1/Parkin function *in vitro* and in cell culture models, understanding the consequences of this dysfunction on neuronal demise must be studied *in vivo*, in the complex milieu of organismal biology. This has been severely hampered by the lack of robust phenotypes in *Pink1/Prkn* knockout mice. In contrast, *Drosophila* models have provided substantial insights in this realm as fly *Pink1/parkin* mutants exhibit extensive disruption of the neuromuscular system presenting, amongst other phenotypes, profound deficits in locomotor behaviors, apoptotic degeneration of flight muscles, progressive degeneration of dopaminergic neurons, all accompanied by morphological and functional breakdown of mitochondria. Consequently, genetic studies using the fly models, primarily using suppression or enhancement of the mutant phenotypes as a sensitive readout, have elucidated several important and conserved features of PINK1/Parkin biology^21^.

Recent studies have shed new light on the *in vivo* role of PINK1/Parkin in vertebrates, and the context in which loss of *Pink1/Prkn* can reveal pathogenic phenotypes. First, combining *Prkn* knockout mutants with a mtDNA mutator strain selectively led to degeneration of nigral dopaminergic neurons, decline in motor ability and increased mitochondrial dysfunction^9^. Extending these observations, Sliter et al.^10^ revealed that this *Prkn^−/−^*;*mutator* combination (or *Pink1^−/−^*;*mutator*) provoked an aberrant innate immune response mediated by the STING pathway, suggesting that the systemic inflammatory response ultimately caused the dopaminergic neurodegeneration and motor deficits. Indeed, genetic loss of *STING* was sufficient to completely prevent the inflammation, motor defect and neurodegeneration in the *Prkn^−/−^*;*mutator* mice. These findings established the STING pathway and, more broadly, aberrant innate immune signaling, as a pathogenic cause and a highly attractive therapeutic target. Moreover, additional work has also implicated *Pink1/Prkn* mutations in inducing aberrant inflammation, albeit via adaptive immunity^22^. However, while the PINK1/Parkin pathway is clearly an ancient mechanism regulating mitochondrial quality control, our data indicate that Sting does not appear to be a fundamental, conserved feature of PINK1/Parkin biology.

The question arises why loss of *Sting* does not suppress *Pink1/parkin* phenotypes in flies when it is capable of completely preventing pathology in mice? At this stage, the answer is unknown and rather puzzling given that innate immune signaling is dysregulated in *parkin* mutants^15^, and Sting performs an analogous function in flies as it does in vertebrates^11^. One possibility is that the aberrant innate immune activation observed in *parkin* mutant flies is not mediated by the presence of cytosolic DNA and activation of the Sting pathway. Currently, there is little if any direct evidence to support this. Moreover, investigating whether induction of mtDNA mutations is required to trigger the innate immune response, as indicated by Sliter et al., our data show that even in the presence of a mtDNA mutator, the Sting immune cascade did not contribute to the neuromuscular phenotypes caused by loss of *Pink1/parkin* in flies. An alternative interpretation is that the *Pink1/parkin* phenotypes are not due to aberrant immune signaling and this may be an epiphenomenon. Supporting this view, many studies have established that loss of Pink1/parkin in flies causes catastrophic mitochondrial disruptions, triggering cell-autonomous apoptosis^16–18^.

Considering this, it isn’t clear from current data why either exhaustive exercise or increased mtDNA mutations should trigger an innate immune response that is mitigated by Pink1/Parkin in mice. The involvement of STING implicates the presence of cytosolic DNA as a trigger. The evidence from Sliter et al. suggests that exhaustive exercise and/or mtDNA mutations is sufficient to induce mitophagy, which if not properly executed by Pink1/Parkin leads to the release of mtDNA and activation of STING signaling. However, it remains unclear how these mitochondrial stresses in the absence of *Pink1/Prkn* lead to release of mtDNA – presumably by loss of integrity and rupture of the mitochondrial boundary membranes. The observed increase in mitophagy in mouse cardiac muscle upon exhaustive exercise is again intriguing as this tissue shares striking structural and functional homology with *Drosophila* flight muscle, further increasing the puzzle as to why the role of Sting does not appear to be a conserved feature of Pink1/parkin biology in flies. Clearly, further work is necessary in order to fully understand the mechanisms linking mitochondrial disruption and immune activation across species.

## Methods

### *Drosophila* stocks and husbandry

Flies were raised under standard conditions in a humidified, temperature-controlled incubator with a 12h:12h light:dark cycle at 25°C, on food consisting of agar, cornmeal, molasses, propionic acid and yeast. Transgene expression was induced using the ubiquitous *da-GAL4* driver. The following strains were obtained from the Bloomington *Drosophila* Stock Center (RRID:SCR_006457): *w*^1118^ (RRID:BDSC_6326), *da-GAL4* (RRID:BDSC_55850), *Sting*^TRiP^ (RRID:BDSC_31565), *Relish*^TRiP^ (RRID:BDSC_33661), *Relish*^E20^ (RRID:BDSC_9457), *UAS-mito-HA-GFP* (RRID:BDSC_8443); and the Vienna Drosophila Resource Center (RRID:SCR_013805): *Sting*^GD^ (P{GD1905}v4031), *Relish*^GD^ (P{GD1199}v49413), and *lacZ* RNAi (P{GD936}v51446) used as a control RNAi. Other lines were kindly provided as follows: *Sting*^ΔRG5^ from A. Goodman^11^, *Pink1*^B9^ mutants from J. Chung^18^, and the *park*^25^ mutants and *UAS-mito-APOBEC1* have been described previously^17,20^. All experiments were conducted using male flies.

### Locomotor assays

The startle induced negative geotaxis (climbing) assay was performed using a counter-current apparatus. Briefly, 20-23 males were placed into the first chamber, tapped to the bottom, and given 10 s to climb a 10 cm distance. This procedure was repeated five times (five chambers), and the number of flies that has remained into each chamber counted. The weighted performance of several group of flies for each genotype was normalized to the maximum possible score and expressed as *Climbing index*^17^.

### Immunohistochemistry and sample preparation

For immunostaining, adult flight muscles were dissected in PBS and fixed in 4% formaldehyde (pH 7.0) for 30 min, permeabilized in 0.3% Triton X-100 for 30 min, and blocked with 0.3% Triton X-100 plus 1% bovine serum albumin in PBS for 1 h at RT. Tissues were incubated with ATP5A antibody (Abcam Cat# ab14748, RRID:AB_301447; 1:500), diluted in 0.3% Triton X-100 plus 1% bovine serum albumin in PBS overnight at 4°C, then rinsed 3 times 10 min with 0.3% Triton X-100 in PBS, and incubated with the appropriate fluorescent secondary antibodies overnight at 4°C. The tissues were washed 2 times in PBS and mounted on slides using Prolong Diamond Antifade mounting medium (Thermo Fischer Scientific).

### Microscopy

Fluorescence imaging was conducted using a Zeiss LSM 880 confocal microscope (Carl Zeiss MicroImaging) equipped with Nikon Plan-Apochromat 100x/1.4 NA oil immersion objectives. Images were prepared using Fiji software (Fiji, RRID:SCR_002285). For thoracic indentations, images were acquired using a Leica DFC490 camera mounted on a Leica MZ6 stereomicroscope.

### Statistical analysis

For behavioral analyses, Kruskal-Wallis non-parametric test with Dunn’s post-hoc correction for multiple comparisons was used. Analyses were performed using GraphPad Prism 8 software (RRID:SCR_002798).

### Data availability

All data that support the findings of this study are available on reasonable request to the corresponding author. The contributing authors declare that all relevant data are included in the paper.

## Acknowledgements

This work is supported by MRC core funding (MC_UU_00015/4, MC-A070-5PSB0 and MC_UU_00015/6) and ERC Starting grant (DYNAMITO; 309742). J.J.L. was supported by an MRC PhD Studentship awarded via the MRC MBU, and S.A. was supported by an MRC Career Development Fellowship. Stocks were obtained from the Bloomington *Drosophila* Stock Center which is supported by grant NIH P40OD018537. We also thank Alan Goodman (Washington State University) for kindly providing the *Sting* mutant line. We thank other members of the Whitworth lab for discussions.

## Author Contributions

J.J.L. designed and performed experiments, and analysed data, with assistance from S.A. A.J.W. conceived the study, designed experiments, analysed the data and supervised the work. A.J.W. wrote the manuscript with input from all authors.

## References

1 Pickrell, A. M. & Youle, R. J. The roles of PINK1, parkin, and mitochondrial fidelity in Parkinson’s disease. Neuron 85, 257–273, doi:10.1016/j.neuron.2014.12.007 (2015).

2 Yamano, K., Matsuda, N. & Tanaka, K. The ubiquitin signal and autophagy: an orchestrated dance leading to mitochondrial degradation. EMBO Rep 17, 300–316, doi:10.15252/embr.201541486 (2016).

3 Cummins, N. & Gotz, J. Shedding light on mitophagy in neurons: what is the evidence for PINK1/Parkin mitophagy in vivo? Cell Mol Life Sci 75, 1151–1162, doi:10.1007/s00018-017-2692-9 (2017).

4 Rodger, C. E., McWilliams, T. G. & Ganley, I. G. Mammalian mitophagy - from in vitro molecules to in vivo models. FEBS J 285, 1185–1202, doi:10.1111/febs.14336 (2017).

5 Whitworth, A. J. & Pallanck, L. J. PINK1/Parkin mitophagy and neurodegeneration-what do we really know in vivo? Curr Opin Genet Dev 44, 47–53, doi:10.1016/j.gde.2017.01.016 (2017).

6 Vincow, E. S. et al. The PINK1-Parkin pathway promotes both mitophagy and selective respiratory chain turnover in vivo. Proceedings of the National Academy of Sciences of the United States of America 110, 6400–6405, doi:10.1073/pnas.1221132110 (2013).

7 Lee, Y., Dawson, V. L. & Dawson, T. M. Animal models of Parkinson’s disease: vertebrate genetics. Cold Spring Harb Perspect Med 2, doi:10.1101/cshperspect.a009324 (2012).

8 Perez, F. A. & Palmiter, R. D. Parkin-deficient mice are not a robust model of parkinsonism. Proceedings of the National Academy of Sciences of the United States of America 102, 2174–2179, doi:10.1073/pnas.0409598102 (2005).

9 Pickrell, A. M. et al. Endogenous Parkin Preserves Dopaminergic Substantia Nigral Neurons following Mitochondrial DNA Mutagenic Stress. Neuron 87, 371–381, doi:10.1016/j.neuron.2015.06.034 (2015).

10 Sliter, D. A. et al. Parkin and PINK1 mitigate STING-induced inflammation. Nature 561, 258–262, doi:10.1038/s41586-018-0448-9 (2018).

11 Martin, M., Hiroyasu, A., Guzman, R. M., Roberts, S. A. & Goodman, A. G. Analysis of Drosophila STING Reveals an Evolutionarily Conserved Antimicrobial Function. Cell Rep 23, 3537–3550 e3536, doi:10.1016/j.celrep.2018.05.029 (2018).

12 Goto, A. et al. The Kinase IKKbeta Regulates a STING- and NF-kappaB-Dependent Antiviral Response Pathway in Drosophila. Immunity 49, 225–234 e224, doi:10.1016/j.immuni.2018.07.013 (2018).

13 Liu, Y. et al. Inflammation-Induced, STING-Dependent Autophagy Restricts Zika Virus Infection in the Drosophila Brain. Cell Host Microbe 24, 57–68 e53, doi:10.1016/j.chom.2018.05.022 (2018).

14 Kounatidis, I. et al. NF-kappaB Immunity in the Brain Determines Fly Lifespan in Healthy Aging and Age-Related Neurodegeneration. Cell Rep 19, 836–848, doi:10.1016/j.celrep.2017.04.007 (2017).

15 Greene, J. C., Whitworth, A. J., Andrews, L. A., Parker, T. J. & Pallanck, L. J. Genetic and genomic studies of Drosophila parkin mutants implicate oxidative stress and innate immune responses in pathogenesis. Hum Mol Genet 14, 799–811, doi:10.1093/hmg/ddi074 (2005).

16 Clark, I. E. et al. Drosophila pink1 is required for mitochondrial function and interacts genetically with parkin. Nature 441, 1162–1166, doi:10.1038/nature04779 (2006).

17 Greene, J. C. et al. Mitochondrial pathology and apoptotic muscle degeneration in Drosophila parkin mutants. Proceedings of the National Academy of Sciences of the United States of America 100, 4078–4083, doi:10.1073/pnas.0737556100 (2003).

18 Park, J. et al. Mitochondrial dysfunction in Drosophila PINK1 mutants is complemented by parkin. Nature 441, 1157–1161, doi:10.1038/nature04788 (2006).

19 Whitworth, A. J. et al. Increased glutathione S-transferase activity rescues dopaminergic neuron loss in a Drosophila model of Parkinson’s disease. Proc Natl Acad Sci USA 102, 8024–8029, doi:10.1073/pnas.0501078102 (2005).

20 Andreazza, S. et al. Mitochondrially-targeted APOBEC1 is a potent mtDNA mutator affecting mitochondrial function and organismal fitness in Drosophila. Nat Commun 10, 3280, doi:10.1038/s41467-019-10857-y (2019).

21 Hewitt, V. L. & Whitworth, A. J. Mechanisms of Parkinson’s Disease: Lessons from Drosophila. Curr Top Dev Biol 121, 173–200, doi:10.1016/bs.ctdb.2016.07.005 (2017).

22 Matheoud, D. et al. Intestinal infection triggers Parkinson’s disease-like symptoms in Pink1(-/-) mice. Nature 571, 565–569, doi:10.1038/s41586-019-1405-y (2019).

